# T-bet marks contact-capable regulatory T cells embedded within Th1–dendritic cell networks in skin-draining lymph nodes

**DOI:** 10.64898/2026.10.05.756696

**Authors:** Amanda H. Priddy, Mitra P. Maz, Olena Kamenyeva, Juraj Kabat, Ethan M. Shevach, Munir Akkaya, Billur Akkaya

## Abstract

Regulatory T cells (Tregs) adapt to inflammatory environments while preserving Foxp3-dependent suppressive identity. In type 1 inflammation, Tregs can upregulate T-bet, but whether T-bet⁺ Tregs represent a stable regulatory subset or exhibit plasticity toward an effector-like state remains incompletely understood. To probe the positioning and interactions of T-bet⁺ Tregs during type 1 inflammation, we performed live lymph node section microscopy in T-bet–ZsGreen Foxp3-RFP dual-reporter mice during a subcutaneous complete Freund’s adjuvant (CFA) challenge. T-bet⁺ Tregs accumulated rapidly in the draining lymph nodes within 24 hours, coincident with the appearance of T-bet⁺ dendritic cells (DCs) and Th1-like CD4⁺ T cells. Analysis of lymph node sections revealed that T-bet⁺ Tregs were incorporated into Th1–DC contact networks, displaying increased contact frequency and contact volume compared with T-bet⁻ Tregs. In parallel, flow cytometric analysis of the immune response showed that T-bet⁺ Tregs maintained a stable regulatory profile distinct from T-bet⁺ effector T cells, with retained high expression of Foxp3 and regulatory-associated surface molecules including FR4, CD25, Nrp-1, CD39, and GARP. They also expressed elevated adhesion and activation-associated molecules, including LFA-1, ICOS, PD-1, CD44, and CD103 compared with T-bet⁻ Tregs, and were enriched for CD103⁺ cells across the inflammatory time course. Adoptive transfer studies further showed that pre-existing T-bet expression enhanced recovery of antigen-specific Tregs from CFA-draining lymph nodes in a context-dependent manner, whereas T-bet expression did not confer a unique advantage in canonical in vitro suppression of Th1 proliferation. Together, these findings identify T-bet⁺ Tregs as a contact-capable regulatory subset embedded within early Th1–DC networks during the priming phase of type 1 inflammation rather than a destabilized or effector-converted Treg population.

## Introduction

Regulatory T cells (Tregs) are central mediators of peripheral tolerance. Their lineage identity and suppressive function are governed by the transcription factor Foxp3, yet Tregs are not phenotypically static^1–3^. Both in the steady state and during inflammation, Tregs acquire context-specific adaptations that allow them to function within distinct immune environments while preserving their core regulatory program^4–8^. One such adaptation is the expression of T-bet, a transcription factor classically associated with type 1 immune responses^6, 8, 9^.

T-bet is induced in multiple immune populations during type 1 inflammation, including CD4⁺ and CD8⁺ effector T cells, natural killer cells, DCs, innate lymphoid cells, inflammatory myeloid populations, and B cells^9–14^ In conventional T cells, antigen presenting cell (APC)-derived inflammatory cues, including IL-12, induce or reinforce T-bet expression, promote Th1 differentiation, and support IFN-γ production^9, 10, 15^. In Foxp3⁺ Tregs, however, the role of T-bet is more complex. Previous studies have shown that T-bet expression in Tregs is required for effective control of type 1 inflammation, in part through induction of CXCR3 and positioning along CXCL9/CXCL10-rich inflammatory gradients^6, 8^. On the other hand, T-bet expression can also be seen in the context of Treg destabilization and effector conversion, where Foxp3 expression is weakened or lost and effector-associated programs emerge^16, 17^. However, this phenomenon appears most prominent in disease states with high inflammatory cytokine load, whereas Foxp3⁺ Tregs are generally stable under many physiologic and inflammatory conditions^17, 18^. With approximately 10–20% of stable Foxp3⁺ Tregs displaying some level of detectable T-bet expression in naïve mice^7, 15^, it is important to understand how T-bet complements the behavior and function of bona fide Tregs within intact lymphoid tissue during type 1 inflammation.

Many experimental models of immunization and organ-specific autoimmunity begin with peripheral delivery of antigen emulsified in CFA, including models such as experimental autoimmune encephalomyelitis, experimental autoimmune uveitis, collagen-induced arthritis, and experimental autoimmune neuritis^19–22^. In such settings, antigen-specific immune responses are initiated in draining lymph nodes before propagating to target tissues^23, 24^. Thus, understanding how T-bet⁺ Tregs behave during CFA-driven type 1 inflammation may provide insight into the regulation of type 1 immune responses in animal models of organ specific autoimmunity^25^.

Here, using live lymph node section microscopy in T-bet–ZsGreen Foxp3-RFP dual-reporter mice, we found that T-bet⁺ Tregs accumulate rapidly within 24 hours of CFA challenge and are highly embedded within Th1–DC contact networks. Flow cytometry analysis demonstrated that T-bet⁺ Tregs retained a stable regulatory phenotype enriched for adhesion, activation, and contact-associated molecules. Although adoptive transfer experiments suggested that pre-existing T-bet expression favors accumulation of antigen-specific Tregs in CFA-draining lymph nodes in vivo, T-bet expression did not confer a unique advantage in canonical in vitro suppression of Th1 proliferation. Together, these findings identify T-bet⁺ Tregs as a contact-capable regulatory subset embedded within early Th1–DC networks, rather than a destabilized or effector-converted Treg population in skin-draining lymph nodes.

## Results

### T-bet⁺ Tregs rapidly accumulate in CFA-draining lymph nodes during early type 1 inflammation

To determine how T-bet-expressing Tregs are positioned during the development of type 1 inflammation, we injected CFA into the flanks of T-bet–ZsGreen Foxp3-RFP reporter mice subcutaneously and analyzed draining inguinal lymph nodes at baseline and at days 1, 4, and 7 after challenge (**Figure 1a**). Live lymph node microscopy followed by histocytometry allowed us to map Th1-like CD4⁺Foxp3⁻T-bet⁺ cells, CD11c⁺ dendritic cells, CD11c⁺T-bet⁺ dendritic cells, and Foxp3⁺T-bet⁺ Tregs within the same tissue volume (**Figure 1a,b**). CFA challenge induced a rapid reorganization of the draining lymph node. By day 1, T-bet⁺ CD4 effector cells and T-bet⁺ dendritic cells were increased, and Foxp3⁺T-bet⁺ Tregs accumulated prominently within the same inflammatory regions (**Figure 1b,c**). Quantification showed that the absolute number of T-bet⁺ Tregs peaked within 24 hours and then declined toward baseline by days 4 and 7 (**Figure 1d**). This increase was not explained simply by proportional expansion of the total Treg compartment: both the frequency of T-bet⁺ cells among Tregs and the absolute number of T-bet⁺ Tregs increased early after challenge (**Figure 1d**).

**Figure 1.**
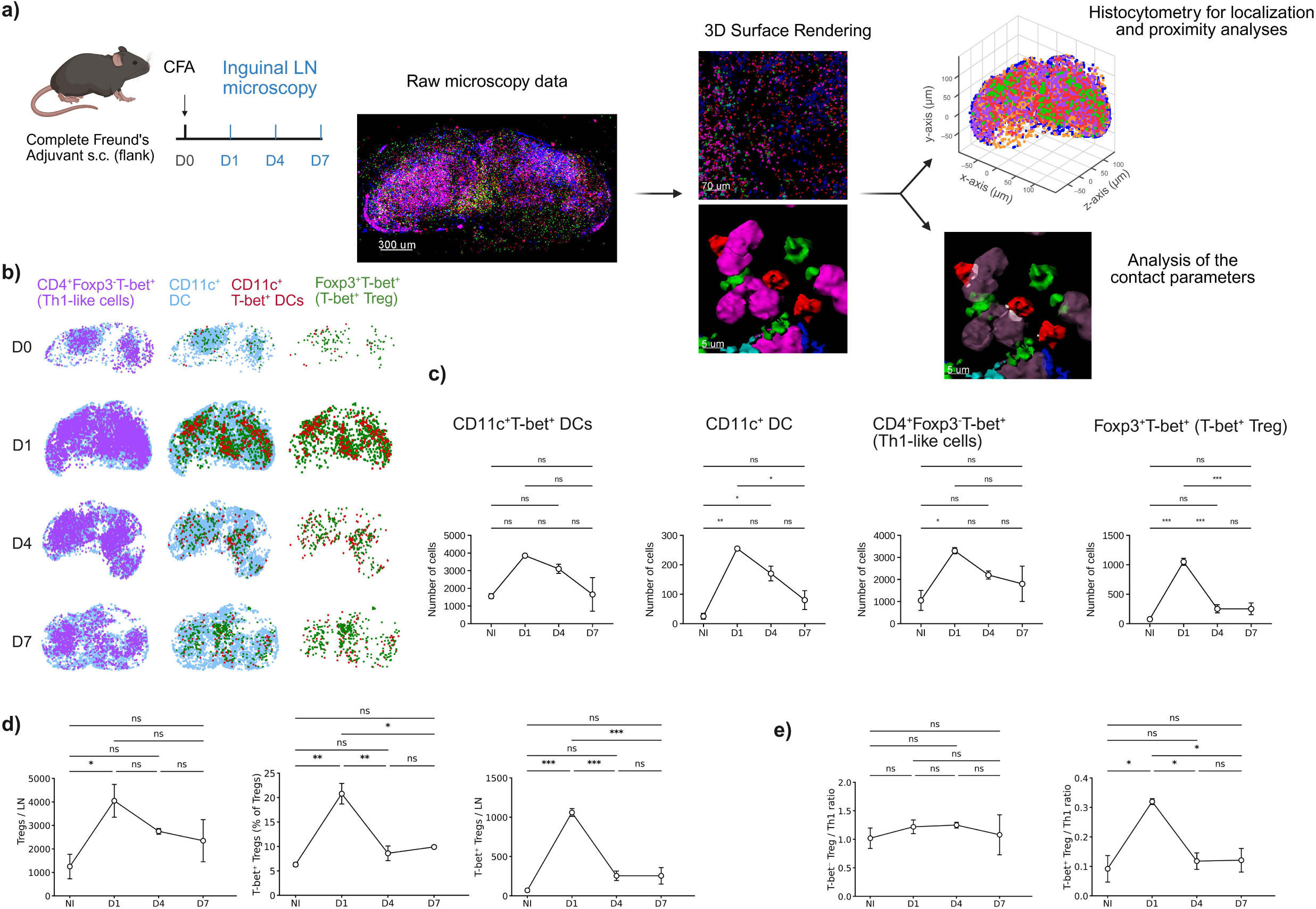
T-bet⁺ Tregs rapidly accumulate with T-bet⁺ dendritic cells and Th1-like cells in CFA-draining inguinal lymph nodes. T-bet–ZsGreen Foxp3-RFP reporter mice were challenged subcutaneously with CFA and draining inguinal lymph nodes were analyzed at baseline and at days 1, 4, and 7 after challenge. **a**, Experimental and imaging workflow. Draining inguinal lymph nodes were harvested, sectioned, labeled with anti-CD4 and anti-CD11c antibodies, and analyzed by live tissue-section confocal microscopy, three-dimensional surface rendering, histocytometry, and contact analysis. Representative raw microscopy data, three-dimensional renderings, coordinate-based histocytometry plots, and contact-analysis outputs are shown. **b**, Histocytometric localization maps generated from Imaris-exported x, y, and z coordinates of individual segmented cells and plotted in FlowJo. Maps show the distribution of CD4⁺Foxp3⁻T-bet⁺ Th1-like cells, CD11c⁺ dendritic cells, CD11c⁺T-bet⁺ dendritic cells, and Foxp3⁺T-bet⁺ Tregs in draining inguinal lymph node sections at the indicated time points. **c**, Quantification of CD11c⁺T-bet⁺ dendritic cells, total CD11c⁺ dendritic cells, CD4⁺Foxp3⁻T-bet⁺ Th1-like cells, and Foxp3⁺T-bet⁺ Tregs per lymph node section. **d**, Quantification of total Tregs per lymph node, percentage of Tregs expressing T-bet, and number of T-bet⁺ Tregs per lymph node. **e**, Ratio of T-bet⁻ Tregs to Th1-like cells and ratio of T-bet⁺ Tregs to Th1-like cells over the inflammatory time course. Graphs show mean ± SEM from n = 3 mice per time point. P values were calculated using one-way ANOVA with Tukey’s multiple comparisons test. ns, not significant; *P < 0.05; **P < 0.01; ***P < 0.001; ****P < 0.0001.

The relationship between Tregs and Th1-like cells was also altered during this early response. Whereas the ratio of T-bet⁻ Tregs to Th1-like cells remained relatively stable, the ratio of T-bet⁺ Tregs to Th1-like cells increased sharply at day 1 before returning toward baseline (**Figure 1e**). Thus, CFA challenge induces a transient early enrichment of T-bet⁺ Tregs in the draining lymph node, coincident with the emergence of T-bet⁺ dendritic cells and Th1-like CD4 T cells.

Consistent with these changes in cell number, T-bet intensity within the T-bet⁺ Treg compartment was dynamically regulated, increasing early after CFA challenge and declining thereafter, whereas T-bet intensity in Th1-like cells remained comparatively stable over the same time course (**Supplementary Figure S1a**). Time-lapse imaging also showed population-specific changes in cell speed during the CFA response in Th1-like cells, T-bet⁻ Tregs, and T-bet⁺ Tregs (**Supplementary Figure S1b**). These findings indicate that the T-bet⁺ Treg compartment is numerically and transcriptionally responsive during the earliest phase of the draining lymph node response.

### T-bet⁺ Tregs are preferentially incorporated into Th1-DC contact networks

We next asked whether T-bet⁺ Tregs merely accumulated in the same lymph node or whether they were physically incorporated into local Th1-associated cellular neighborhoods. Three-dimensional surface rendering and histocytometry were used to identify Treg–Th1 contacts across the draining lymph node. Compared with T-bet⁻ Tregs, T-bet⁺ Tregs showed a markedly greater frequency of contact with Th1-like CD4⁺Foxp3⁻T-bet⁺ cells (**Figure 2a**). This difference was evident across the time course and was most pronounced during the early inflammatory response. These results indicate that T-bet expression identifies a Treg subset that is highly incorporated into Th1-associated contact networks within the draining lymph node.

**Figure 2.**
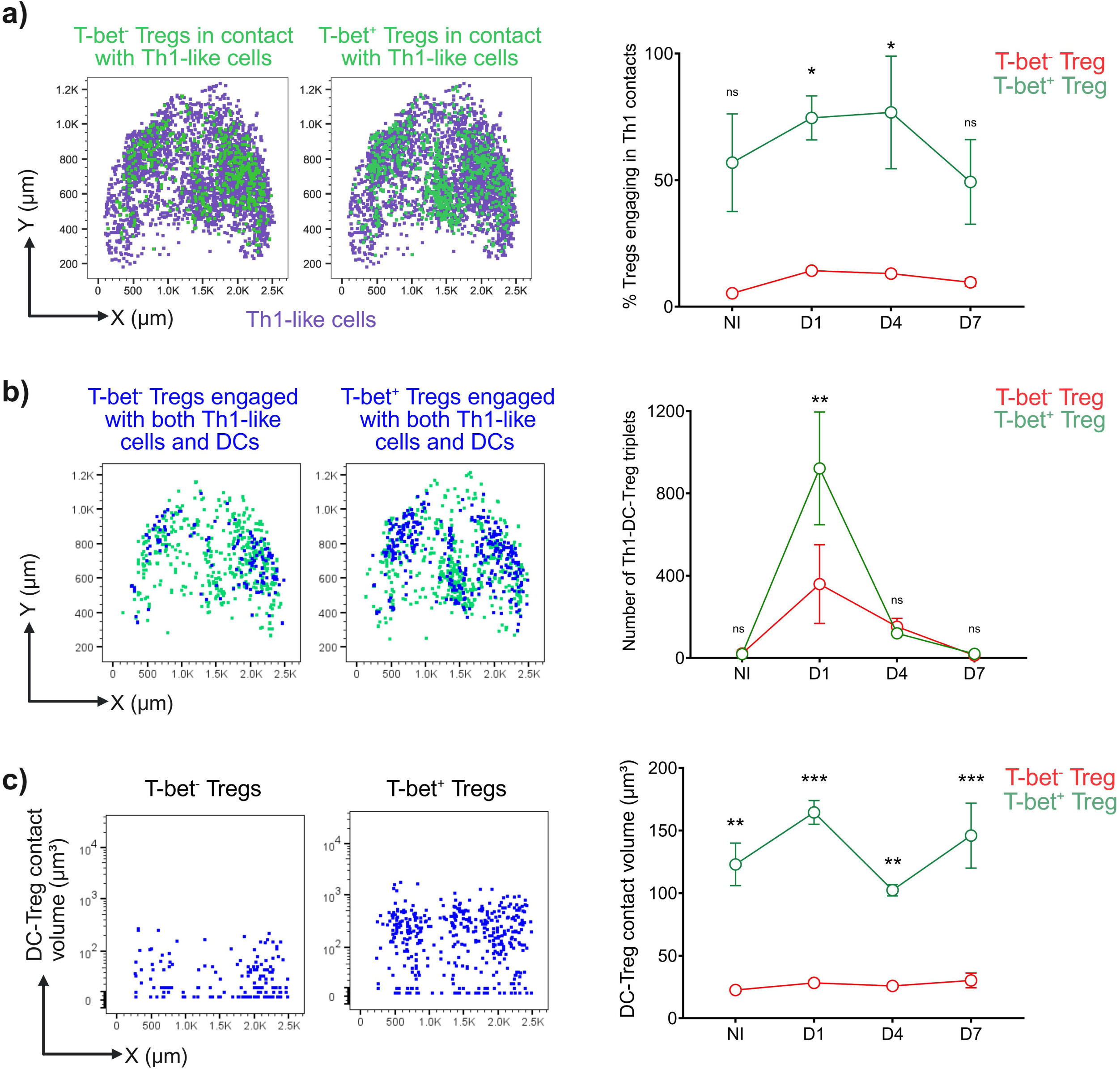
T-bet⁺ Tregs preferentially participate in Th1 contact networks in CFA-draining inguinal lymph nodes. Using the same imaging datasets shown in Figure 1, we analyzed three-dimensional surface-surface colocalization among Th1-like cells, Tregs, and dendritic cells. **a**, Representative coordinate-based maps showing CD4⁺Foxp3⁻T-bet⁺ Th1-like cells and Foxp3⁺ Tregs engaged in Th1-like cell–Treg contacts, separated according to T-bet⁻ or T-bet⁺ Treg status. Th1-like cells are shown in purple, and Tregs in contact with Th1-like cells are shown in green. The graph shows the percentage of T-bet⁻ and T-bet⁺ Tregs engaging in contacts with Th1-like cells at baseline and at days 1, 4, and 7 after CFA challenge. **b**, Representative coordinate-based maps showing the subset of T-bet⁻ and T-bet⁺ Tregs engaged with both Th1-like cells and dendritic cells. Green points indicate Tregs in contact with Th1-like cells, and blue points indicate Tregs within Th1–DC–Treg triplets. The graph shows the number of Th1–DC–Treg triplets across the inflammatory time course. **c**, Representative plots showing DC–Treg contact volume for T-bet⁻ and T-bet⁺ Tregs engaged in Th1–DC–Treg contact networks. Contact volume was calculated from Imaris-derived three-dimensional surface-surface colocalization volume and is shown in μm³. The graph shows DC–Treg contact volume for T-bet⁻ and T-bet⁺ Tregs across the inflammatory time course. Graphs show mean ± SEM from n = 3 mice per time point. P values were calculated using two-way ANOVA with Sidak’s multiple comparisons test for comparisons between T-bet⁻ and T-bet⁺ Tregs at matched time points. ns, not significant; *P < 0.05; **P < 0.01; ***P < 0.001.

Because Th1 responses are organized through interactions with antigen-presenting cells, we next examined whether T-bet⁺ Tregs were associated with dendritic cell-containing contact structures. Histocytometric analysis identified Th1–DC–Treg triplets and allowed comparison of structures involving T-bet⁻ versus T-bet⁺ Tregs (**Figure 2b**). T-bet⁺ Tregs were more frequently associated with Th1–DC contact networks than T-bet⁻ Tregs. At day 1, the number of Th1–DC–Treg triplets increased prominently, and triplets involving T-bet⁺ Tregs were more abundant than those involving T-bet⁻ Tregs (**Figure 2b**). Within these Th1–DC–Treg contact networks, DC–Treg contact volume was also greater for T-bet⁺ Tregs than for T-bet⁻ Tregs (**Figure 2c**). By later time points, these contact structures declined, paralleling the contraction of the T-bet⁺ Treg compartment. Together, these data indicate that T-bet⁺ Tregs are not positioned at the margins of the developing type 1 response. Rather, they are embedded within the inflammatory cellular circuitry itself, occupying Th1–DC contact niches where antigen presentation, effector T cell activation, and regulatory control are likely to converge.

### T-bet⁺ Tregs retain regulatory identity while acquiring adhesion- and contact-associated features

The spatial enrichment of T-bet⁺ Tregs within Th1–DC contact networks raised the question of whether these cells represent stable regulatory cells or a plastic effector-like population. To address this, we performed flow-cytometric phenotyping of T-bet⁻ Teff, T-bet⁺ Teff, T-bet⁻ Tregs, and T-bet⁺ Tregs across the CFA time course using a broad panel of regulatory, activation, trafficking, adhesion, and lineage-associated markers. Principal-component analysis of the markers that most clearly distinguished these populations showed that the four populations occupied distinct phenotypic spaces (**Figure 3a**). T-bet⁺ Tregs remained clearly separated from T-bet⁺ Teff cells, indicating that T-bet expression did not simply impose a shared Th1-like phenotype across lineages. Notably, the T-bet⁺ Treg compartment remained relatively stable across baseline, day 1, day 4, and day 7, despite the marked early increase and later contraction in T-bet⁺ Treg number observed by histocytometry (**Figure 3a**).

**Figure 3.**
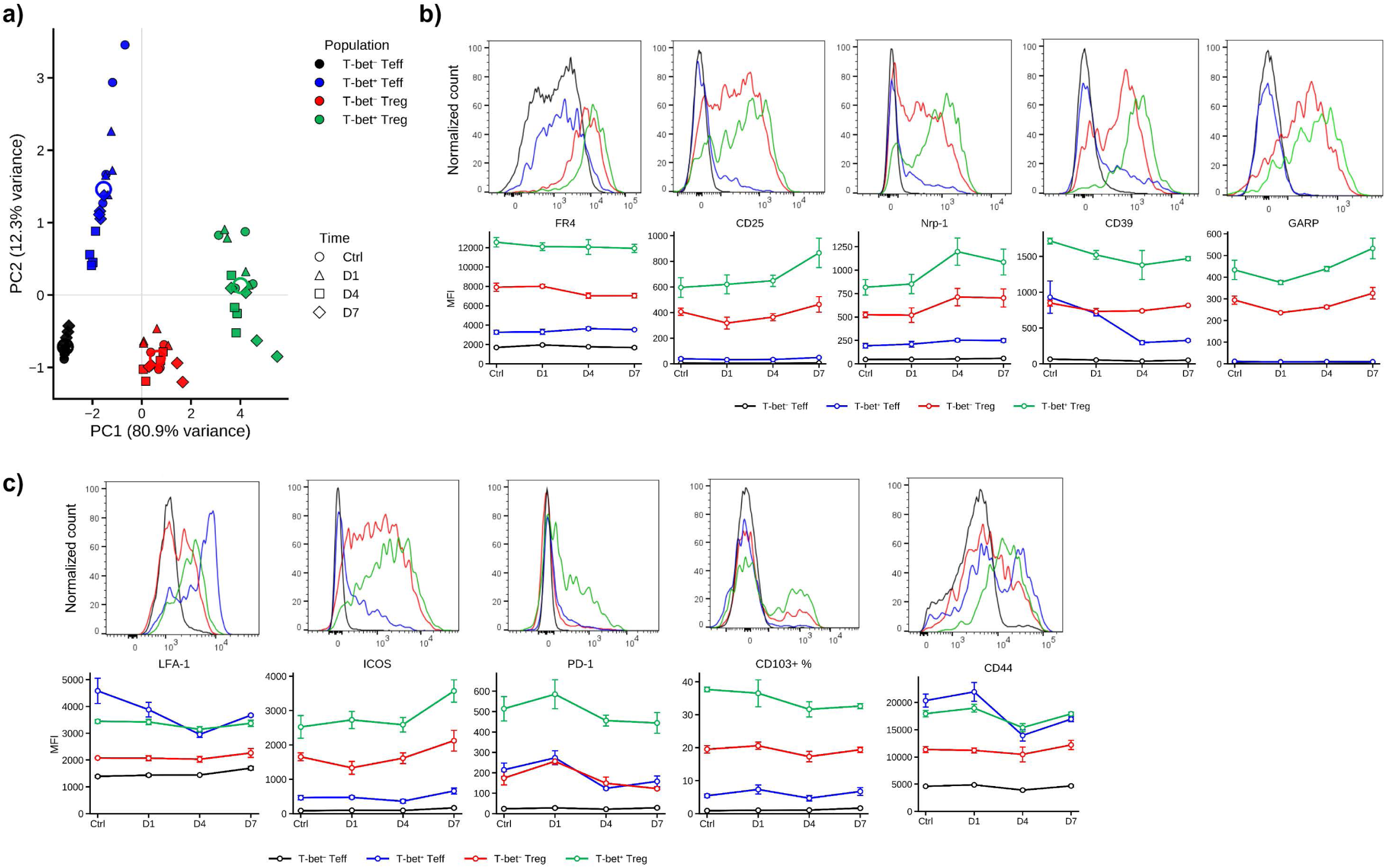
T-bet⁺ Tregs retain a stable regulatory phenotype distinct from T-bet⁺ effector T cells during CFA-driven inflammation. Draining inguinal lymph nodes from T-bet–ZsGreen Foxp3-RFP reporter mice were analyzed by flow cytometry at baseline and after CFA challenge. Live singlet CD4⁺ cells were divided into T-bet⁻Foxp3⁻ conventional CD4⁺ T cells, T-bet⁺Foxp3⁻ Th1-like cells, T-bet⁻Foxp3⁺ Tregs, and T-bet⁺Foxp3⁺ Tregs based on endogenous T-bet–ZsGreen and Foxp3-RFP reporter expression. **a**, Principal component analysis of flow-cytometric marker expression in the four indicated CD4⁺ T cell populations across the inflammatory time course. Symbols denote time point, and colors denote population. **b**, Representative histograms and time-course quantification of regulatory-associated markers FR4, CD25, Nrp-1, CD39, and GARP in the indicated populations. **c**, Representative histograms and time-course quantification of adhesion, activation, and contact-associated markers LFA-1, ICOS, PD-1, CD103, and CD44. T-bet⁺ Tregs remained phenotypically distinct from T-bet⁺ effector T cells and maintained elevated expression of regulatory and contact-associated markers despite dynamic changes in their abundance. Graphs show mean ± SEM from n = 2–4 mice per time point. Data are representative of two independent experiments.

T-bet⁺ Tregs retained high expression of regulatory-associated molecules, including FR4, CD25, Nrp-1, CD39, and GARP (**Figure 3b**). These markers were consistently enriched in T-bet⁺ Tregs compared with T-bet⁻ Tregs and distinguished T-bet⁺ Tregs from T-bet⁺ Teff cells, supporting the interpretation that T-bet⁺ Tregs preserve a canonical regulatory phenotype. In parallel, T-bet⁺ Tregs expressed elevated levels of molecules associated with activation, adhesion, and cellular interaction, including LFA-1, ICOS, PD-1, CD103, and CD44 (**Figure 3c**). Additional markers examined, including CTLA-4, did not show clear differences between T-bet⁺ and T-bet⁻ Tregs and were therefore not included in the main phenotypic summary. CD103 positivity was particularly enriched within the T-bet⁺ Treg compartment. Whereas only a small fraction of Teff cells were CD103⁺, a substantially larger fraction of T-bet⁺ Tregs expressed CD103 across the time course (**Figure 3c**). Thus, T-bet⁺ Tregs are not simply characterized by higher CD103 fluorescence intensity; rather, the T-bet⁺ Treg compartment is compositionally enriched for CD103⁺ cells. These data indicate that T-bet⁺ Tregs are phenotypically stable regulatory cells with an adhesion- and contact-associated profile, consistent with their localization within Th1–DC-rich tissue niches.

### T-bet deficiency does not abolish antigen-driven Treg expansion but reduces CXCR3 and CD39 expression

We next tested whether T-bet was required for antigen-specific Treg accumulation in vivo. OT-II Tregs from T-bet-sufficient and T-bet-deficient backgrounds were co-transferred into recipient mice, followed by subcutaneous challenge with OVA-CFA or PBS in separate recipient mice (**Figure 4a**). Draining inguinal lymph nodes were analyzed seven days later using flow cytometry (**Figure 4a,b; Supplementary Figure S3a**). Both T-bet-sufficient and T-bet-deficient OT-II Tregs proliferated in response to OVA-CFA and were recovered from antigen-draining lymph nodes (**Figure 4c–e**). T-bet deficiency did not eliminate antigen-driven expansion or recovery of transferred Tregs in this setting (**Figure 4d,e**). Spleen analysis from the same experiment similarly showed recovery and proliferation of transferred OT-II Tregs without a genotype-specific defect in the T-bet-deficient population (**Supplementary Figure S3c**). However, T-bet-deficient Tregs expressed lower levels of CXCR3 and CD39 compared with T-bet-sufficient Tregs (**Figure 4f**), whereas CD73 expression was not significantly altered (**Supplementary Figure S3b**). Thus, although T-bet was not absolutely required for antigen-driven Treg proliferation or lymph node recovery, it supported expression of molecules associated with type 1 positioning and regulatory function. These findings suggest that T-bet contributes to the inflammatory adaptation of antigen-specific Tregs without being strictly required for their proliferation in response to antigen plus CFA.

**Figure 4.**
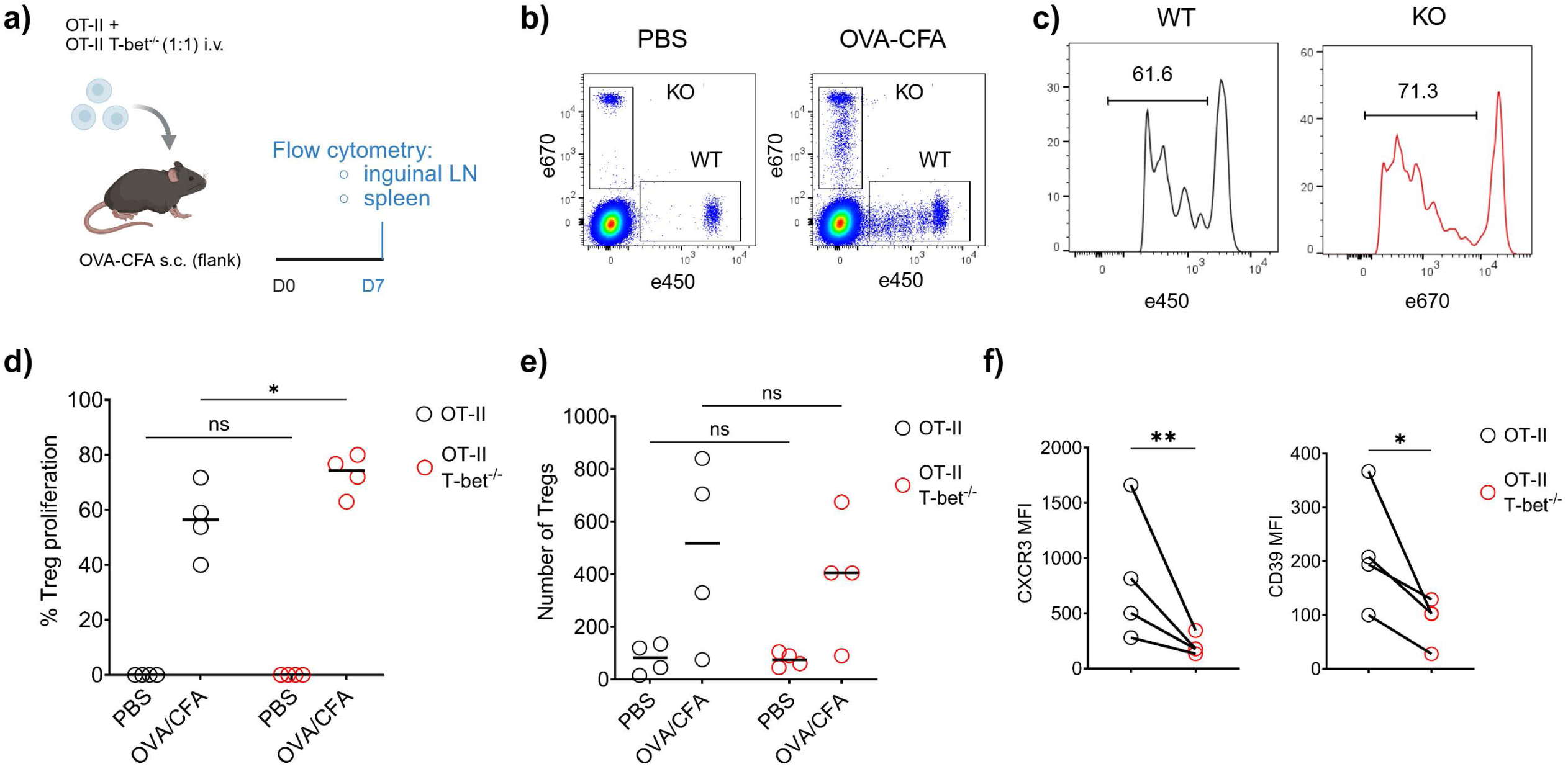
T-bet is dispensable for antigen-driven OT-II Treg proliferation and recovery but supports CXCR3 and CD39 expression. Neutral OT-II Tregs and T-bet-deficient OT-II Tregs were generated in vitro, differentially labeled with e450 or e670 proliferation dye, mixed at a 1:1 ratio, and transferred intravenously into recipient mice challenged with OVA_323-339_-CFA or PBS. Draining inguinal lymph nodes were analyzed by flow cytometry at day 7 after challenge. **a**, Experimental design. **b**, Representative flow cytometry plots showing recovery of transferred wild-type and T-bet-deficient OT-II Tregs from PBS- and OVA-CFA-draining inguinal lymph nodes. **c**, Representative proliferation-dye histograms showing proliferation of recovered wild-type and T-bet-deficient OT-II Tregs. **d**, Quantification of Treg proliferation in PBS- and OVA-CFA-draining inguinal lymph nodes. **e**, Quantification of recovered transferred OT-II Tregs. **f**, Paired comparison of CXCR3 and CD39 expression in recovered wild-type and T-bet-deficient OT-II Tregs from OVA-CFA-draining inguinal lymph nodes. Data are representative of two independent experiments with n = 2–4 mice per group, as indicated. P values in **d** and **e** were calculated using two-way ANOVA with Sidak’s multiple comparisons test. P values in **f** were calculated using paired two-tailed Student’s t-test. ns, not significant; *P < 0.05; **P < 0.01.

### Pre-existing T-bet expression enhances recovery of antigen-specific Tregs from CFA-draining lymph nodes

We next asked whether a pre-existing T-bet-associated program could enhance Treg accumulation within inflamed draining lymph nodes. To test this, we generated OT-II iTregs under neutral or T-bet-polarizing conditions and transferred them into recipient mice challenged with OVA-CFA (**Figure 5a**). Because these experiments relied on in vitro-generated T-bet⁺ iTregs as donor cells, we first compared their type 1-associated phenotype with endogenous T-bet⁺ thymic/natural Tregs from T-bet–ZsGreen Foxp3-RFP reporter mice (**Supplementary Figure S2a**). T-bet⁺ iTregs and endogenous T-bet⁺ tTregs showed broadly similar CXCR3 staining and IL-12Rβ1 positivity, while IL-12Rβ2 expression remained low or absent in both populations (**Supplementary Figure S2a**). In a separate comparison, neutral iTregs, T-bet-deficient iTregs, and T-bet⁺ iTregs were restimulated with PMA/ionomycin and assessed for IFN-γ and IL-2 by intracellular cytokine staining (**Supplementary Figure S2b**). T-bet⁺ iTregs did not show an effector-like cytokine profile under these pharmacologic restimulation conditions. Thus, although in vitro-generated T-bet⁺ iTregs are not identical to endogenous T-bet⁺ tTregs, they preserved key features of the T-bet-associated Treg phenotype relevant to type 1 inflammatory adaptation.

**Figure 5.**
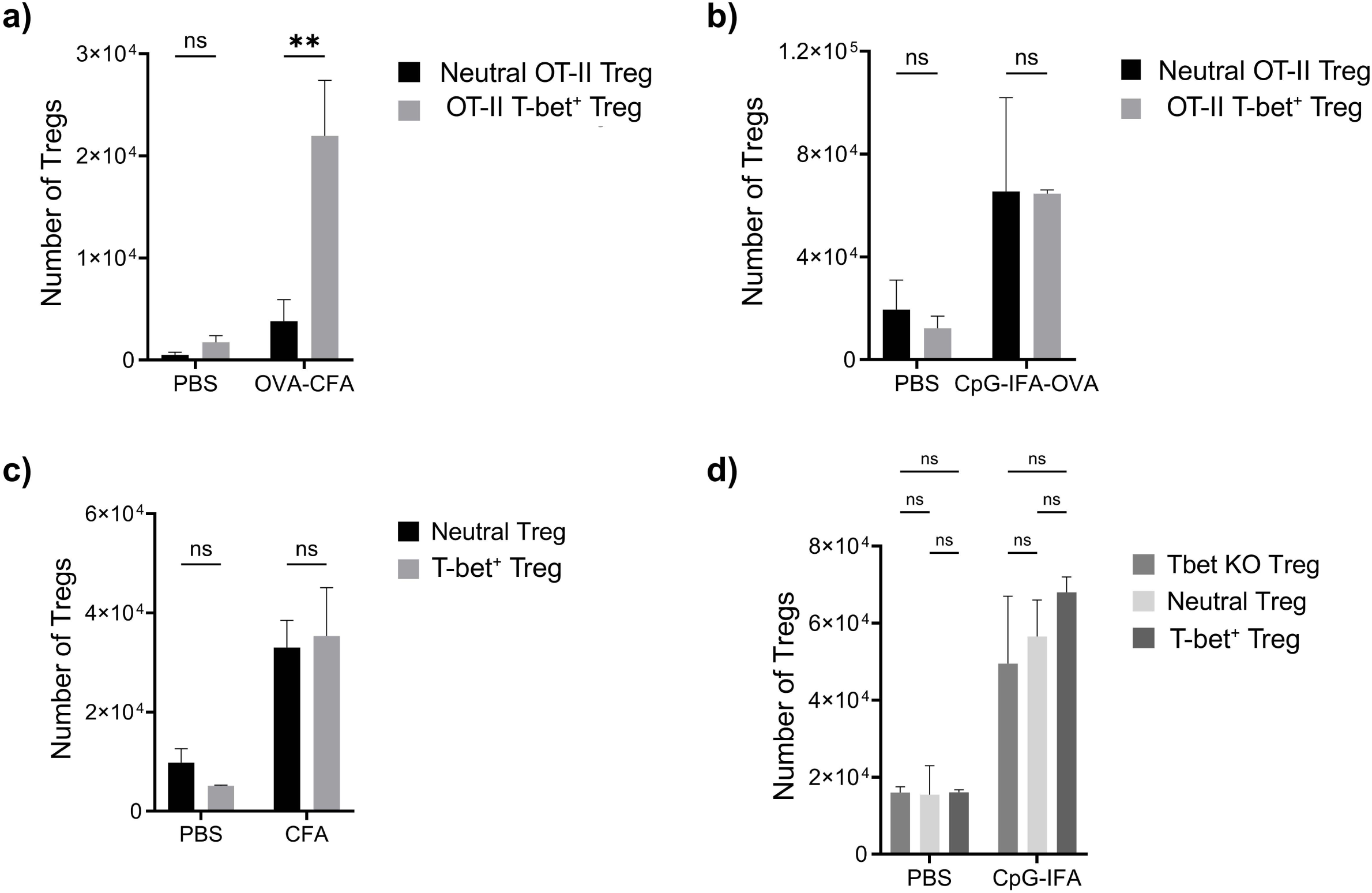
Pre-existing T-bet expression enhances recovery of antigen-specific Tregs from CFA-draining inguinal lymph nodes. Neutral and T-bet-polarized Tregs were generated in vitro, labeled with e450 or e670, and transferred i.v. into recipient mice challenged with the indicated stimuli. Draining inguinal lymph nodes were analyzed by flow cytometry. **a,** Quantification of recovered neutral OT-II Tregs and T-bet-polarized OT-II Tregs from PBS- or OVA_323-339_-CFA-draining inguinal lymph nodes. **b**, Quantification of recovered neutral OT-II Tregs and T-bet-polarized OT-II Tregs from PBS- or OVA_323-339_-CpG-IFA-draining inguinal lymph nodes. Under CpG-IFA conditions, T-bet polarization did not confer a recovery advantage. **c**, Quantification of recovered polyclonal neutral and T-bet-polarized Tregs from PBS- or CFA-draining inguinal lymph nodes. **d**, Quantification of recovered T-bet-deficient, neutral, and T-bet-polarized polyclonal Tregs from PBS- or CpG-IFA-draining inguinal lymph nodes. Graphs show mean ± SEM. Data are representative of two independent experiments with n = 2–4 mice per group, as indicated. P values in a–c were calculated using two-way ANOVA with Sidak’s multiple comparisons test. P values in d were calculated using two-way ANOVA with Tukey’s multiple comparisons test. ns, not significant; *P < 0.05; **P < 0.01.

Compared with neutral OT-II Tregs, T-bet-polarized OT-II Tregs were recovered in greater numbers from OVA-CFA-draining lymph nodes (**Figure 5a**). This advantage was context-dependent. In lymph nodes draining OVA emulsified in CpG-IFA, neutral and T-bet-polarized OT-II Tregs were recovered at similar levels (**Figure 5b**). Likewise, when polyclonal neutral and T-bet-polarized Tregs were transferred, T-bet polarization did not confer a clear recovery advantage in CFA-draining lymph nodes (**Figure 5c**). Finally, in CpG-IFA-draining lymph nodes, T-bet-deficient, neutral, and T-bet-polarized polyclonal Tregs were recovered at similar levels (**Figure 5d**). These findings suggest that T-bet expression is most beneficial when paired with antigen specificity in a CFA-driven type 1 inflammatory environment.

Because these cells were delivered intravenously, these experiments do not distinguish increased recruitment from blood, increased retention, local survival, or proliferation within the draining lymph node. However, they support the conclusion that a pre-existing T-bet-associated program can enhance accumulation of antigen-specific Tregs in the lymph node responding to CFA-associated inflammation. Together with the endogenous imaging and phenotyping data, these results suggest that T-bet is not simply a marker acquired after arrival in an inflammatory site, but may also identify a regulatory program that improves Treg participation in antigen-driven type 1 immune niches.

### T-bet expression does not confer a unique advantage in canonical in vitro suppression of Th1 proliferation

Finally, we asked whether T-bet expression directly altered the ability of Tregs to suppress Th1 proliferation in vitro. Th1 cells were stimulated in the absence of Tregs or co-cultured with T-bet-deficient Tregs, T-bet⁻ neutral Tregs, or T-bet⁺ Tregs (**Figure 6a**). All Treg populations suppressed Th1 proliferation relative to cultures without Tregs (**Figure 6b**). T-bet⁺ Tregs did not show a unique advantage over T-bet⁻ or T-bet-deficient Tregs in this canonical suppression assay (**Figure 6b**). In contrast to their effect on proliferation, none of the Treg populations substantially reduced PMA/ionomycin-induced IFN-γ production by Th1 cells (**Supplementary Figure S4a,b**). Thus, under maximal pharmacologic restimulation conditions, Tregs did not measurably impair the cytokine-producing capacity of Th1 cells, and T-bet expression did not confer a unique advantage in suppressing this response. Instead, the combined imaging, phenotyping, and adoptive-transfer data support a model in which T-bet identifies Tregs with the positioning and inflammatory adaptation needed to participate in local Th1/DC contact networks during early type 1 inflammation.

**Figure 6.**
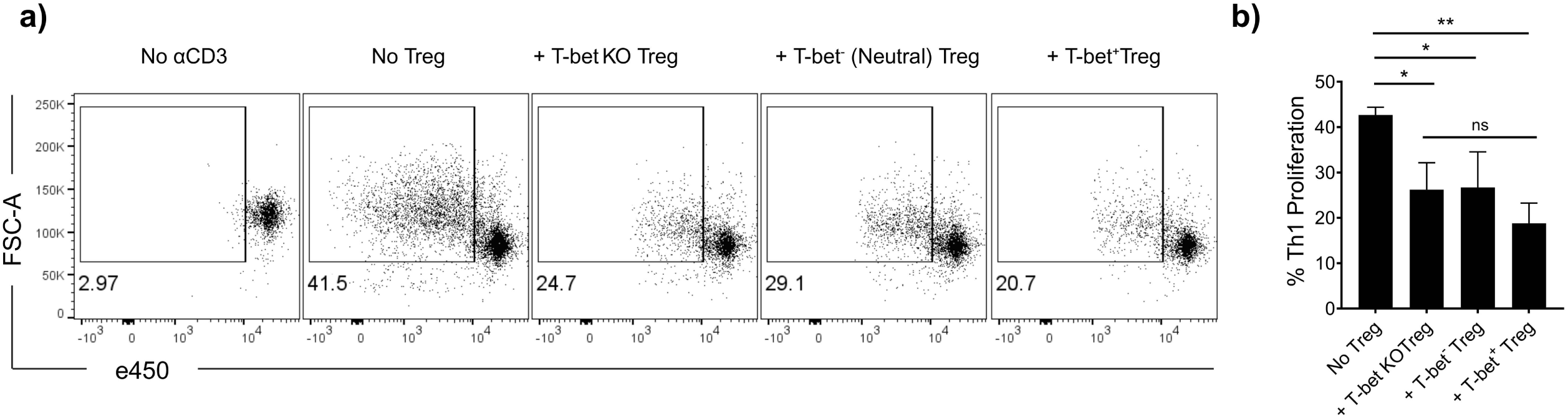
T-bet expression does not confer a unique advantage in canonical in vitro suppression of Th1 proliferation. **a-b**, Th1-polarized responder cells were cultured in the absence or presence of Tregs generated under the indicated conditions, and responder-cell proliferation was assessed by flow cytometry. **a**, Representative flow cytometry plots showing Th1 proliferation in the absence of anti-CD3 stimulation, in the absence of Tregs, or in the presence of T-bet-deficient Tregs, neutral T-bet⁻ Tregs, or T-bet⁺ Tregs. **b**, Quantification of Th1 proliferation under the indicated conditions. Graphs show mean ± SEM from the indicated biological replicates. P values were calculated using one-way ANOVA with Tukey’s multiple comparisons test. ns, not significant; *P < 0.05; **P < 0.01.

## Discussion

Tregs must adapt to inflammatory environments while preserving their Foxp3-dependent regulatory identity to prevent immunopathology during ongoing immune responses^1, 2^. In type 1 inflammation, this adaptation includes expression of T-bet, a transcription factor classically associated with Th1 differentiation^4, 6, 9^. T-bet has been shown to promote CXCR3 expression in Tregs and to support their accumulation at sites of Th1-mediated inflammation, establishing the concept that Tregs co-opt inflammatory transcriptional programs to regulate matching effector responses^6^. Subsequent work further demonstrated that T-bet expression in Tregs can represent a stable regulatory state rather than a transient step toward effector conversion, and that T-bet⁺ Tregs can selectively restrain Th1 and CD8⁺ T cell activation^7, 8^. Our findings extend this framework by showing that, in skin-draining lymph nodes, T-bet identifies Tregs that rapidly enter the cellular networks of type 1 inflammation.

A central finding of this study is that T-bet⁺ Tregs are numerically dynamic but phenotypically stable. T-bet⁺ Tregs accumulated rapidly within 24 hours after CFA challenge and then declined toward baseline at later time points. However, PCA and marker-level analyses showed that these cells remained phenotypically distinct from T-bet⁺ effector CD4⁺ T cells. T-bet⁺ Tregs retained Foxp3 expression and expressed higher levels of regulatory-associated molecules, including FR4, CD25, Nrp-1, CD39, and GARP, compared with T-bet⁻ Tregs. They also expressed increased LFA-1, ICOS, PD-1, CD44, and CD103, consistent with activation, APC experience, adhesion, and niche engagement. The CD103 data were particularly informative because the T-bet⁺ Treg compartment was enriched for CD103⁺ cells, rather than merely showing a shift in CD103 MFI. These data do not determine whether the early increase in T-bet⁺ Tregs reflects recruitment of pre-existing T-bet⁺ Tregs from blood, entry through afferent lymphatics from inflamed skin, or local induction of T-bet in Tregs already present within the lymph node. Nevertheless, they support the interpretation that T-bet⁺ Tregs are not unstable intermediates drifting toward an effector-like state, but bona fide regulatory cells with an adhesion- and contact-associated phenotype.

The spatial organization of these cells provides the major conceptual advance of this study. T-bet⁺ Tregs were not simply increased in number; they were specifically incorporated into Th1–DC contact networks. They showed increased contact frequency with Th1-like cells, increased representation within Th1–DC–Treg triplets, and greater DC–Treg contact volume within these networks compared with T-bet⁻ Tregs. Thus, T-bet⁺ Tregs were not positioned at the margins of the developing type 1 response. Rather, they were embedded within the inflammatory cellular circuitry itself, occupying cellular niches where DCs, Th1-like cells, and Tregs converge. This interpretation is consistent with imaging studies showing that Tregs regulate immune priming through spatially organized contacts with dendritic cells. Intravital and explanted lymph node imaging studies demonstrated that Tregs can limit productive interactions between conventional CD4⁺ T cells and antigen-bearing dendritic cells during priming^26, 27^. Our previous work further showed that antigen-specific Tregs form strong interactions with dendritic cells and deplete cognate peptide–MHC class II complexes from the DC surface, selectively limiting presentation of the relevant antigen while preserving bystander access to the same DCs^28^. Together, these studies support a model in which Tregs do not simply suppress through global effects but can act locally within antigen-presenting microenvironments.

Recent work in tumor-draining lymph nodes and tumors provides an especially relevant parallel. CXCR3⁺ Tregs have been shown to interact preferentially with CXCL9-producing BATF3⁺ type 1 dendritic cells, thereby limiting antigen cross-presentation and CD8⁺ T cell responses^29^. In lung-cancer-draining mediastinal lymph nodes, tissue-specific IFN-γ abundance promoted a Th1-like effector Treg program characterized by T-bet and CXCR3 expression, close interaction with DC1s, and MHC-II-dependent suppression of cytotoxic T cell priming^30^. Our findings extend this concept to CFA-driven inflammation, where T-bet⁺ Tregs were similarly positioned within DC–effector T cell networks in draining lymph nodes. The adoptive-transfer experiments add an in vivo recovery correlate to this spatial pattern. Pre-existing T-bet expression enhanced recovery of antigen-specific Tregs from OVA-CFA-draining lymph nodes, but this advantage was not reproduced uniformly across OVA-CpG-IFA or polyclonal transfer settings. This distinction is informative because CFA and CpG-IFA can both support type 1-skewing immune responses, but they are not interchangeable adjuvant environments. CFA contains mycobacterial components in an oil emulsion and can induce antigen-specific IFN-γ and IL-17A responses in draining lymph nodes through multiple mycobacterial pattern-recognition pathways including NOD2, Mincle/CARD9, and inflammasome-associated signaling^31–33^. CpG-B ODN 1826, by contrast, provides a more defined TLR9-biased stimulus within an IFA emulsion and can promote IL-12-dependent Th1-skewing responses^34–36^. Thus, the recovery advantage of T-bet-polarized OT-II Tregs in OVA-CFA may not reflect a universal benefit of T-bet expression under all type 1-skewing conditions, but rather a particular fit between antigen recognition, T-bet-associated Treg programming, and the cellular organization generated by CFA. Because the transferred cells were delivered intravenously, these experiments do not distinguish whether increased recovery reflects more efficient entry from blood, enhanced retention, increased survival, or local proliferation within the draining lymph node. Nevertheless, the data are consistent with the idea that T-bet expression is most useful when antigen specificity and CFA-driven type 1 inflammatory cues converge, in line with earlier work linking T-bet to CXCR3-dependent accumulation of Tregs in Th1 inflammatory environments^6^.

Importantly, T-bet expression did not confer a unique advantage in canonical in vitro suppression of Th1 proliferation. T-bet-deficient, neutral, and T-bet⁺ Tregs were all capable of suppressing Th1 proliferation, indicating that T-bet is not required for generic suppressive activity in this reductionist assay. Rather than pointing to increased suppressive potency per cell, these findings suggest that the major contribution of T-bet in this system is to identify Tregs adapted to the anatomical and cellular context in which type 1 immunity is being initiated. In supplementary analyses, Tregs also did not substantially reduce PMA/ionomycin-induced IFN-γ production by Th1 cells, regardless of whether the Tregs were wild-type, T-bet-deficient, or T-bet-polarized. PMA/ionomycin provides strong pharmacologic restimulation that bypasses proximal antigen-receptor-dependent priming events; therefore, this assay primarily tests whether prior Treg co-culture abolishes the cytokine-producing capacity of Th1 cells under maximal restimulation conditions. The preservation of IFN-γ production after PMA/ionomycin does not exclude regulation of Th1 cytokine responses in vivo, where Tregs may act upstream of cytokine induction by limiting antigen presentation, costimulation, IL-2 availability, extracellular ATP metabolism, DC function, or cellular access to APCs^28, 37–40^. Taken together, the strongest differences between T-bet⁺ and T-bet⁻ Tregs emerged in spatial organization, contact-associated phenotype, and in vivo recovery from CFA-draining lymph nodes rather than suppressive potency in vitro. This suggests that the major advantage of T-bet⁺ Tregs may not be stronger suppression per cell, but improved participation in the anatomical site where inflammatory priming occurs.

In summary, our findings identify T-bet⁺ Tregs as a contact-capable regulatory subset embedded within early Th1–DC networks in skin-draining lymph nodes. Rather than marking unstable Tregs or effector conversion, T-bet identifies a phenotypically stable regulatory state whose abundance changes rapidly during type 1 inflammation while its regulatory and adhesion-associated identity is maintained. We propose that T-bet marks Tregs that regulate type 1 inflammation from within the inflammatory niche, not from its periphery.

## Materials and Methods

### Mice

T-bet–ZsGreen reporter mice were provided by J. Zhu, National Institute of Allergy and Infectious Diseases, National Institutes of Health, as NIAID line 8419, and bred in-house. Foxp3-RFP reporter mice, originally generated by the Flavell laboratory, were obtained from The Jackson Laboratory as C57BL/6-Foxp3^tm1Flv/J^, strain 008374. T-bet–ZsGreen mice were crossed to Foxp3-RFP mice to generate T-bet–ZsGreen Foxp3-RFP double-reporter mice. T-bet-deficient mice were obtained from J. Zhu, NIAID, NIH, as NIAID line 8451 and from The Jackson Laboratory as B6.129S6-Tbx21^tm1Glm/J^, strain 004648. OVA_323-339_-specific OT-II TCR-transgenic mice were obtained from The Jackson Laboratory and crossed as needed to generate antigen-specific donor populations. All mouse strains were maintained on a C57BL/6 background. Mice were maintained in The Ohio State University University Laboratory Animal Resources and National Institutes of Health animal facilities in compliance with Institutional Animal Care and Use Committee standards. All animal experiments were conducted in accordance with approved protocols OSU IACUC 2022A00000061 and NIAID LISB-13E.

### Reagents and flow cytometry

Cells were cultured in sterile complete RPMI medium consisting of RPMI 1640 supplemented with 10% heat-inactivated FBS, 50 U/ml penicillin, 50 μM streptomycin, 1 mM sodium pyruvate, 2 mM L-glutamine, 0.1 mM non-essential amino acids, 50 μM 2-mercaptoethanol, and 10 mM HEPES. For magnetic separations, cells were maintained in filtered and degassed MACS buffer consisting of PBS supplemented with 0.5% BSA and 2 mM EDTA. These medium and buffer conditions follow those used in our previous studies. For flow cytometry, single-cell suspensions were prepared from draining inguinal lymph nodes and stained in FACS buffer consisting of PBS or HBSS supplemented with 2% FBS, 1% HEPES, and 10 mM sodium azide. Cells were blocked with anti-CD16/32 antibody, rat serum, and mouse serum before staining. Viability was assessed using a fixable live/dead near-infrared viability dye. Surface staining was performed with fluorochrome-conjugated antibodies at 4°C for 30 min in the dark. Where intracellular staining was required, cells were fixed and permeabilized according to the manufacturer’s instructions. Flow cytometry data were acquired on BD LSRFortessa and analyzed using FlowJo software, consistent with our prior workflow Antibodies were purchased from BioLegend unless otherwise indicated. The following anti-mouse antibody clones were used for data shown in this study: CD45.1 clone A20, CD45.2 clone 104, CD4 clone RM4-5 or GK1.5, Foxp3 clone FLK-16s, CD103 clone 2E7, CD44 clone IM7, CD11c clone N418, GARP clone YGIC86, CD25 clone PC61, ICOS clone 7E.17G9, CD11a/LFA-1 clone H155-78, IFN-γ clone XMG1.2, CD39 clone Duha59, CD304/Neuropilin-1 clone 3E12, FR4 clone 12A5, and CXCR3 clone CXCR3-173. For the comparison of T-bet⁺ thymic Tregs and in vitro-generated T-bet⁺ iTregs, antibodies against IL-12Rβ1/CD212 clone 114 and IL-12Rβ2 clone 305719 were used. For time-course phenotyping, live singlet CD4⁺ cells were divided into four populations based on endogenous T-bet–ZsGreen and Foxp3-RFP reporter expression: T-bet⁻ Foxp3⁻ conventional CD4⁺ T cells, T-bet⁺ Foxp3⁻ Th1-like CD4⁺ T cells, T-bet⁻ Foxp3⁺ Tregs, and T-bet⁺ Foxp3⁺ Tregs. Marker expression was quantified across these populations at baseline and after CFA challenge. For intracellular cytokine analysis, cells were restimulated with The Cell Stimulation Cocktail 500X (Thermofisher) containing phorbol 12-myristate 13-acetate (PMA), ionomycin, brefeldin A and monensin prior to staining for IFN-γ.

### CFA, CpG-IFA immunization, and lymph node harvest

To induce type 1 inflammation in skin-draining lymph nodes, mice were injected subcutaneously in one or both flanks with CFA (Sigma-Aldrich), OVA_323-339_ peptide emulsified in CFA, or OVA_323-339_ peptide emulsified with CpG-B ODN 1826 in incomplete Freund’s adjuvant (IFA), as indicated. For CFA-containing antigen immunizations, mice received 100 μg OVA_323-339_ peptide in 100 μl emulsion per flank. For CpG-IFA immunizations, mice received 100 μg OVA _323–339_ peptide and 50 μg CpG-B ODN 1826 (InvivoGen) emulsified in IFA in a final volume of 100 μl per flank. Draining inguinal lymph nodes were harvested at the indicated time points after immunization. Control samples were obtained from separate PBS-injected mice. For time-course phenotyping experiments, draining inguinal lymph nodes were collected from control mice and from mice at days 1, 2, 4, and 7 after CFA challenge. Single-cell suspensions were prepared for flow cytometric analysis, or intact lymph nodes were processed for live tissue-section imaging as described below.

### Live inguinal lymph node section imaging

Inguinal lymph nodes from T-bet–ZsGreen Foxp3-RFP reporter mice were harvested and kept on ice in PBS supplemented with 1% BSA. Lymph nodes were embedded in prewarmed RPMI containing 2% agarose and chilled to allow agarose polymerization. Residual fat and connective tissue were removed using a Leica MZ6 StereoZoom microscope (Leica Microsystems). Lymph nodes were suspended in 38°C 2% agarose in RPMI and kept over ice. Following complete agarose solidification, complete RPMI was added, and blocks were cut into 200 μm sections with a Leica VT1000 S vibrating blade microtome (Leica Microsystems) at speed 5 in ice-cold PBS. Prior to imaging, complete RPMI medium was added to tissue sections, and sections were incubated at 37°C, 5% CO₂ for 2 h.

eFluor 450-conjugated anti-CD11c antibody clone N418 and Alexa Fluor 647-conjugated anti-CD4 antibody clone RM4-5 were added at a 1:50 dilution in complete RPMI. Sections were held down with tissue anchors (Warner Instruments) in 35-mm dishes with a 14-mm glass coverslip bottom (MatTek) and imaged with a Leica SP8 inverted five-channel confocal microscope equipped with an environmental chamber (NIH, Division of Scientific Equipment and Instrumentation Services) and a motorized stage. The microscope configuration was set up for four-dimensional analysis (x, y, z, t) of cell organization and contact dynamics within tissue sections. A diode laser for 405 nm, an argon laser for 488 and 514 nm, a diode-pumped solid-state laser for 561 nm, and a HeNe laser for 594 and 633 nm excitation wavelengths were tuned to minimal power, between 0.3% and 2%. Z-stacks of 10–25 μm were collected. Mosaic images of whole lymph nodes were generated by acquiring multiple z-stacks using a motorized stage to cover the whole lymph node area and assembling them into tiled images with LAS X software. For time-lapse contact analysis, tiled z-stacks were collected over 1–4 h.

### Histocytometry and contact-volume analyses

Analysis was performed using Imaris v10.0.0 (Bitplane) as previously described^41^. Briefly, image stacks were stabilized and deconvolved in Huygens where appropriate before import into Imaris for three-dimensional surface reconstruction for each cell population based on fluorescence intensity thresholds and surface-to-surface contact analysis. Endogenous ZsGreen signal was used to identify T-bet-expressing cells, endogenous RFP signal was used to identify Foxp3⁺ Tregs, CD11c staining was used to identify dendritic cells, and CD4 staining was used to identify CD4⁺ T cell populations. This approach allowed classification of CD11c⁺T-bet⁻ dendritic cells, CD11c⁺T-bet⁺ dendritic cells, CD4⁺Foxp3⁻T-bet⁺ Th1-like cells, CD4⁺Foxp3⁺T-bet⁻ Tregs, and CD4⁺Foxp3⁺T-bet⁺ Tregs within intact inguinal lymph node sections. For spatial localization analyses, the x, y, and z coordinates of each segmented surface object were exported from Imaris in micrometers, together with fluorescence-intensity and population-identity parameters. Exported datasets were saved as comma-separated value files and imported into FlowJo for further analysis and visualization. Coordinate-based three-dimensional localization plots were generated in FlowJo by plotting the exported x, y, and z coordinates of individual cells within each lymph node section. For motility analyses, cell tracks were generated from time-lapse image sequences in Imaris, and cell speed was calculated from reconstructed tracks. Cell-cell contact analyses were performed from Imaris-generated three-dimensional surface objects. Surface-surface colocalization parameters were exported for each relevant cell-pair interaction, including contact status and three-dimensional colocalized/contact volume in μm³. These Imaris-derived parameters were exported as comma-separated value files and imported into FlowJo for downstream gating, visualization, and quantification. Th1–Treg contacts were defined as surface-surface colocalization events between CD4⁺Foxp3⁻T-bet⁺ Th1-like cells and Foxp3⁺ Tregs. Th1–DC–Treg triplets were defined as multicellular contact events in which a Th1-like cell, a dendritic cell, and a Treg were simultaneously associated within the same three-dimensional image volume based on surface-surface colocalization parameters. Contact frequency and contact volume were compared between T-bet⁻ and T-bet⁺ Tregs.

### In vitro Treg and Th1 differentiation

For iTreg differentiation, naïve CD4⁺ T cells were isolated from T-bet–ZsGreen Foxp3-RFP mice on either polyclonal or OT-II TCR-transgenic backgrounds. Cells were seeded at 3 × 10⁵ cells per well in 1 ml complete RPMI into 24-well sterile tissue culture plates (Corning) coated with anti-CD3ε and anti-CD28 antibodies (BioLegend), as described^42^. For neutral iTreg differentiation, cells were supplemented with 100 IU/ml recombinant human IL-2 (NCI, Frederick) and 5 ng/ml recombinant human TGF-β (PeproTech) in complete RPMI. For T-bet⁺ iTreg differentiation, 10 ng/ml IFN-γ (PeproTech) and 10 μg/ml anti-IL-4 antibody clone 11B11 (Bio X Cell) were added in addition to IL-2 and TGF-β. Cells were cultured at 37°C, 5% CO₂ for 3–4 days and sorted for live RFP⁺ or ZsGreen⁺RFP⁺ populations for neutral and T-bet⁺ iTregs, respectively, using a BD FACSAria III sorter. For Th1 differentiation, naïve CD4⁺ T cells were seeded into 24-well sterile tissue culture plates coated with 2 μg/ml anti-CD3ε and 1 μg/ml anti-CD28 antibodies. Cells were cultured in complete RPMI supplemented with 10 ng/ml recombinant IL-12 and 10 μg/ml anti-IL-4 antibody clone 11B11. Cells were maintained at 37°C, 5% CO₂ for 3–4 days and sorted as live CD4⁺Foxp3⁻T-bet⁺ cells before use as responder cells in in vitro assays.

### Adoptive transfer

Mice were anesthetized in an induction chamber with 1.5% isoflurane USP (Baxter). A total of 1 × 10⁶ Tregs labeled with e450 or e670 proliferation dye (Thermo Fisher Scientific) were injected intravenously in 100 μl sterile PBS through the retro-orbital sinus. Recipient mice were challenged subcutaneously with OVA_323-339_-CFA, OVA_323-339_-CpG-IFA, CFA, or PBS as indicated. Draining inguinal lymph nodes were harvested at the indicated time points, and transferred cells were quantified by flow cytometry using congenic markers, proliferation dyes, and/or reporter expression.

### Statistical analysis

Statistical analysis was conducted on Prism v10.3.1 (GraphPad). Statistical tests and relevant P values for each figure are indicated in the figure or figure legend.

## Data Availability Statement

Data supporting the findings of this study are available from the corresponding authors upon request. Processed source data and repository information will be added before journal submission.

## Author contributions

M.A. and B.A. jointly supervised the work. B.A. conceived the study, developed the conceptual framework, directed final data interpretation, assembled the final figures, and wrote the manuscript. M.A. contributed to conceptual development, data interpretation, supervision, and critical revision of the manuscript. A.H.H. and M.M. performed experiments, generated data, and contributed to data analysis. O.K. and J.K. contributed to live lymph node imaging, image processing, and histocytometry analysis. E.M.S. provided resources, supervision, and critical intellectual input during the initial experimental phase and critical review of the manuscript. All authors reviewed and approved the final manuscript.

## Acknowledgements

This study was supported by NIH/NIAID extramural grant DP2AI154451, research funds from The Ohio State University College of Medicine, and the Intramural Research Program of the National Institute of Allergy and Infectious Diseases, NIH, under project ZIAAI000224. We thank J. Zhu for providing the T-bet–ZsGreen reporter and T-bet-deficient mice and for helpful discussions throughout the development of this work.

---

**Supplementary Figure 1. T-bet intensity and cell motility are dynamically regulated during CFA-driven inflammation.**

T-bet–ZsGreen Foxp3-RFP reporter mice were challenged subcutaneously with CFA, and draining inguinal lymph node sections were analyzed by live tissue-section imaging at baseline and at days 1, 4, and 7 after challenge. **a**, T-bet reporter intensity was quantified in CD4⁺Foxp3⁻T-bet⁺ Th1-like cells and Foxp3⁺T-bet⁺ Tregs. **b**, Cell speed was quantified from time-lapse imaging in Th1-like cells, T-bet⁻ Tregs, and T-bet⁺ Tregs across the same inflammatory time course. Violin plots show the distribution of cell speeds; red lines indicate median and interquartile range. Data are from n = 3 mice per time point. P values were calculated using one-way ANOVA with Tukey’s multiple comparisons test. ns, not significant; *P < 0.05; **P < 0.01; ***P < 0.001; ****P < 0.0001.

**Supplementary Figure 2. In vitro-generated T-bet⁺ iTregs share key type 1-associated features with endogenous T-bet⁺ thymic/natural Tregs.**

**a**, Endogenous T-bet⁺ tTregs and in vitro-generated T-bet⁺ iTregs were compared by flow cytometry using T-bet–ZsGreen and Foxp3-RFP reporter expression. Representative flow cytometry plots show the gating of T-bet⁺ thymic/natural Tregs and T-bet⁺ iTregs. Histograms show expression of CXCR3, IL-12Rβ1, and IL-12Rβ2 in the indicated populations. T-bet⁻ thymic/natural Tregs are shown as an additional reference population where indicated. **b**, Neutral iTregs, T-bet-deficient iTregs, and T-bet⁺ iTregs were restimulated with PMA/ionomycin under pharmacologic restimulation conditions and analyzed by intracellular cytokine staining. Representative histograms show IFN-γ and IL-2 expression in the indicated iTreg populations.

**Supplementary Figure 3. Gating strategy and spleen analysis for OT-II adoptive-transfer experiments.**

**a,** Representative flow cytometry gating strategy used to identify transferred Tregs in adoptive-transfer experiments. Sequential gates show live cells, lymphocytes, singlets, congenically marked donor-cell populations, and Foxp3⁺ transferred Tregs. **b**, Paired analysis of CD73 expression in recovered wild-type and T-bet-deficient OT-II Tregs from OVA_323-339_-CFA-draining inguinal lymph nodes. **c**, Spleen analysis from the adoptive-transfer experiment shown in Figure 4. Graphs show proliferation and recovery of transferred wild-type and T-bet-deficient OT-II Tregs from spleens of mice receiving PBS or OVA_323-339_-CFA challenge. Data are representative of two independent experiments with n = 2–4 mice per group, as indicated. P value in b was calculated using paired two-tailed Student’s t-test. P values in c were calculated using two-way ANOVA with Sidak’s multiple comparisons test. ns, not significant.

**Supplementary Figure 4. Tregs do not suppress IFN-γ production by Th1 cells after maximal pharmacologic restimulation.**

Th1-polarized responder cells were cultured alone or with Tregs generated under the indicated conditions. Cells were restimulated with PMA/ionomycin in the presence of brefeldin A and monensin before intracellular staining for IFN-γ. **a**, Representative flow cytometry plots showing IFN-γ production by Th1 cells in the absence of anti-CD3 stimulation, in the absence of Tregs, or in the presence of T-bet-deficient Tregs, neutral T-bet⁻ Tregs, or T-bet⁺ Tregs. **b**, Quantification of IFN-γ⁺ Th1 cells under the indicated conditions. Treg co-culture did not significantly reduce IFN-γ production after maximal pharmacologic restimulation, and T-bet expression did not confer a unique effect on this response. Graphs show mean ± SEM from the indicated biological replicates. P values were calculated using one-way ANOVA with Tukey’s multiple comparisons test. ns, not significant.

