## Supplementary figures and images for "T-bet marks contact-capable regulatory T cells embedded within Th1–dendritic cell networks in skin-draining lymph nodes"

### Supplementary Figure 1

#### Supplementary Figure S1

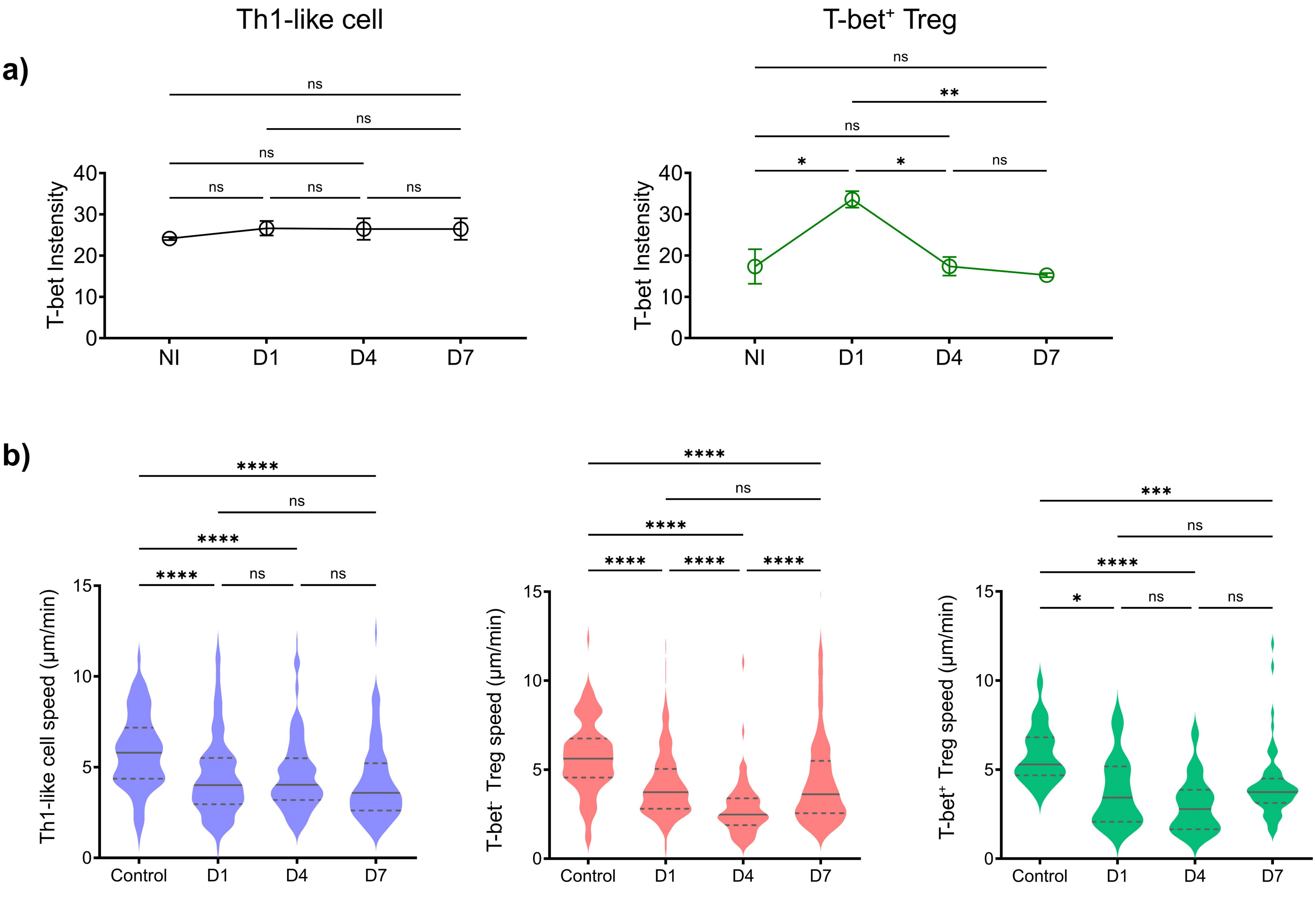

### Supplementary Figure 2

Supplementary Figure S2

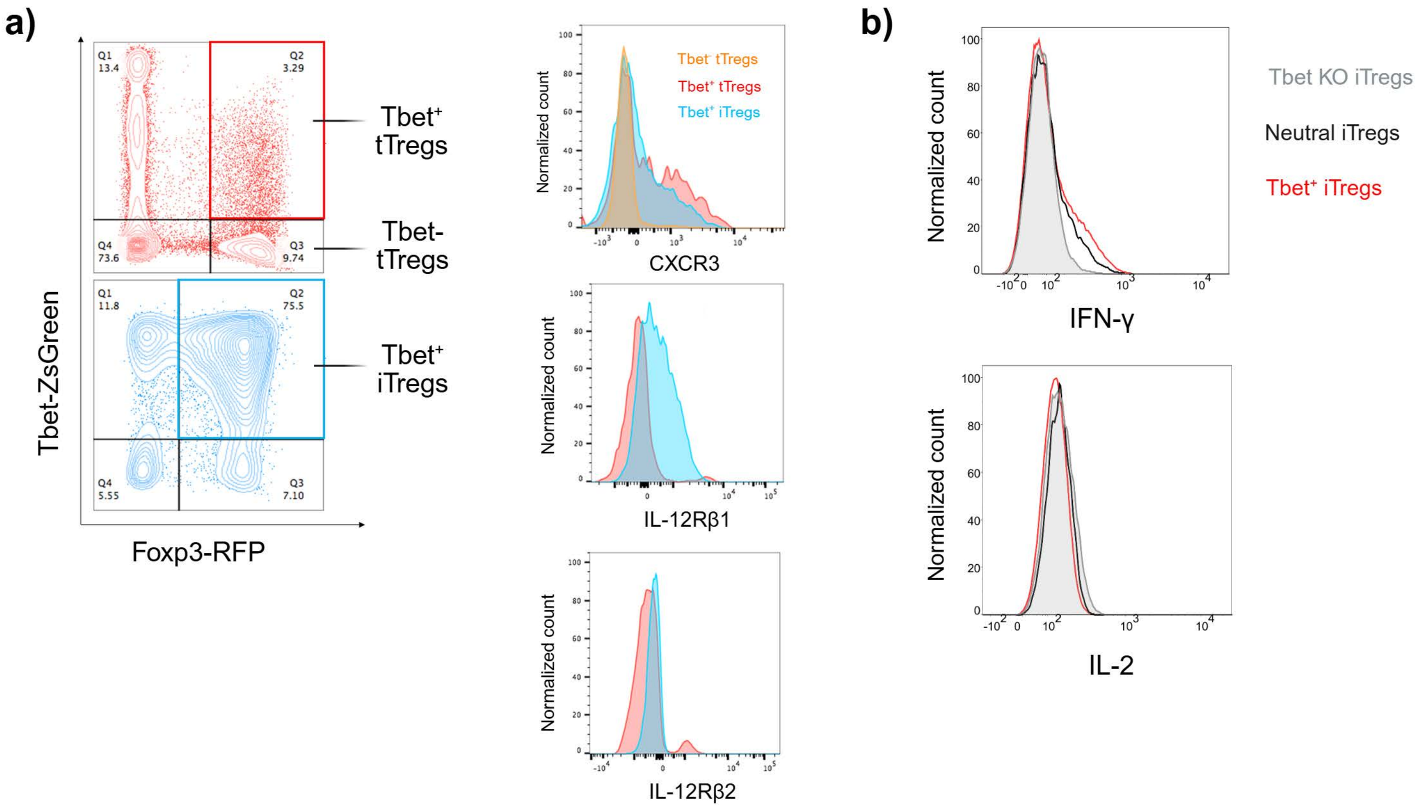

### Supplementary Figure 3

#### Supplementary Figure S3

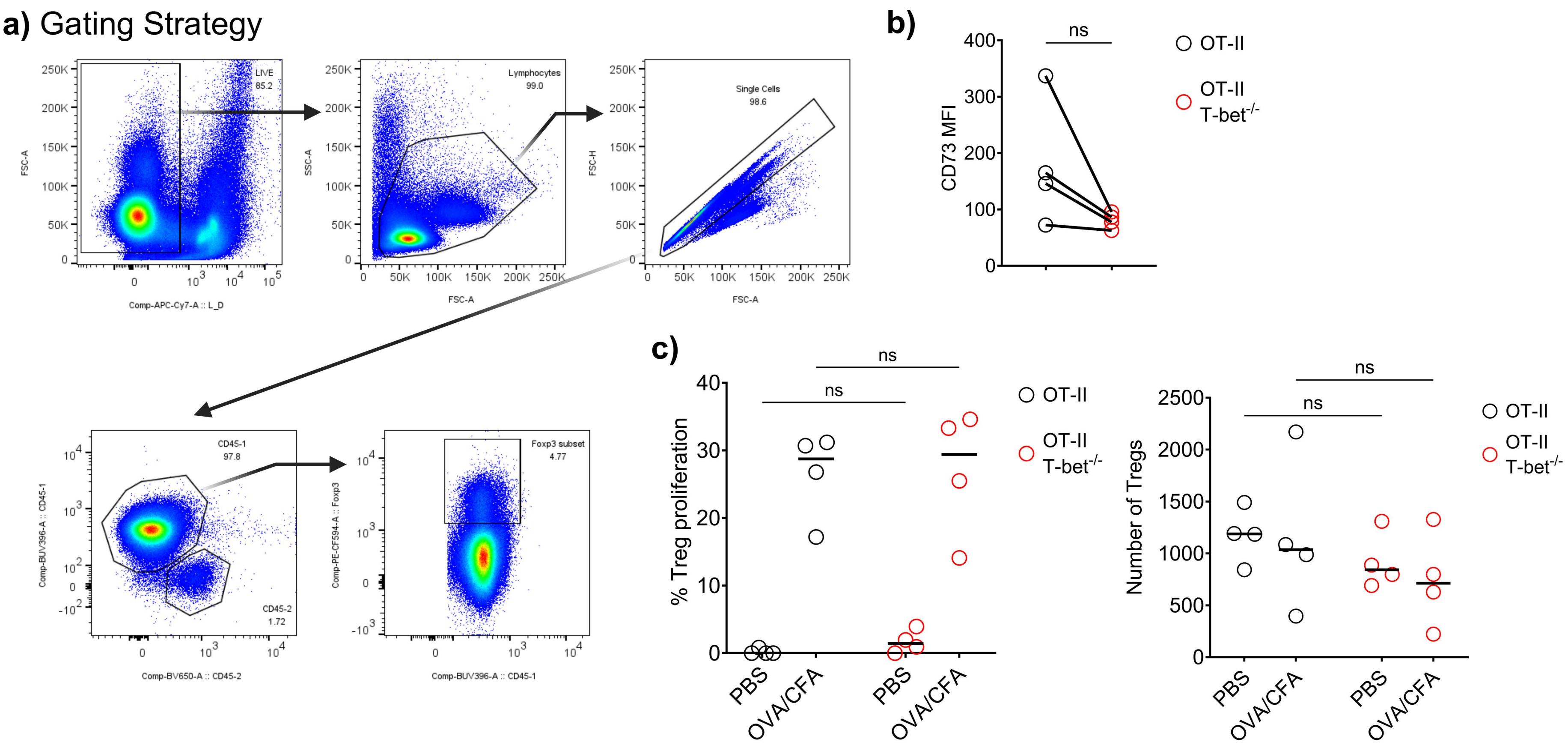

### Supplementary Figure 4

Supplementary Figure S4:

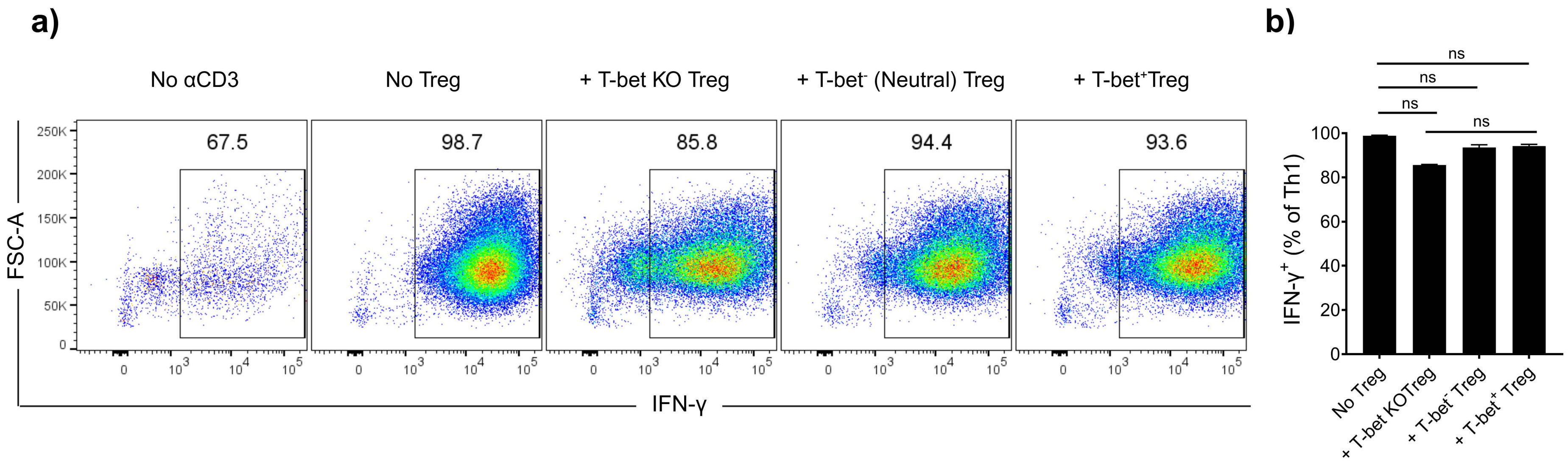
